# Symmetry Analysis and Ancestral Sequence Reconstruction Reveal a Symmetrical Translocation Pathway and Activity Determinants of ZIP Metal Transporter

**DOI:** 10.64898/2026.05.13.724914

**Authors:** Yao Zhang, Tianqi Wang, Hongyan Zhao, Jian Hu

## Abstract

Membrane transporters frequently exhibit internal structural symmetry, reflecting evolutionary origins through gene duplication and fusion events, and this feature has been widely used to infer transport mechanisms. Yet its application to the ZIP (Zrt-/Irt-like protein) family of divalent metal transporters, which are crucial for trace metal homeostasis, has not been fully explored. Here, we apply combined symmetry analysis and ancestral sequence reconstruction to study the ZIP family. We show that internal symmetry is broadly present across prokaryotic ZIPs and is more prominent in reconstructed ancestral sequences, supporting an evolutionary pathway in which the modern 8-transmembrane (TM) ZIP fold arose from duplication, domain rearrangement, and fusion of an ancestral 4-TM protein. Leveraging this symmetry framework, we identify a continuous and symmetric metal translocation pathway composed of symmetric entrance, transport, and exit sites in the ZIP fold, and define gates that ensure alternating access. Application of this structural model to human ZIP4 enables identification of the gate-forming residues, and functional studies reveal that two residues (T529 and V533) in the external gate play a crucial role in controlling metal transport. In addition, comparison with ancestral sequences uncovers a set of LIV-1 subfamily-specific metal chelating residues (D504, E541, and D544) that break the symmetry in the ancestral sequences. Functional studies showed that these residues play distinct roles in transport. Together, our study demonstrates that combining internal symmetry analysis and ancestral sequence reconstruction for the ZIP family facilitates elucidation of metal translocation mechanism and identification of the subfamily-specific features that confer functional specialization.

## Introduction

Membrane transporters frequently exhibit internal structural symmetry, often manifested as inverted repeats, a feature widely thought to arise from early gene duplication and fusion events during evolution (1,2). Such pseudo-symmetry is a hallmark of many transporter families and has proven highly informative for understanding transport mechanisms (3,4). Symmetry analysis has therefore emerged as a powerful method for dissecting transporter function, particularly in cases where dynamic conformational states are difficult to capture experimentally (5,6). A prominent example of internal symmetry in membrane transporters is the Zrt-/Irt-like protein (ZIP) family (7-10), which is ubiquitously expressed across all domains of life and mediates the cellular uptake of essential trace elements, including zinc, iron, manganese, and others. In humans, 14 ZIPs exert critical and diverse physiological functions, and their dysfunction is frequently associated with diseases arising from disrupted metal homeostasis (11-19).

The best structurally characterized ZIP is the prototypical bacterial ZIP from *Bordetella bronchiseptica* (BbZIP) (20-22), which is believed to utilize an elevator-type transport mode to achieve alternating access (21,23,24), even though only inward-facing conformations (IFC) have been experimentally determined. The inverted repeats in BbZIP structure were initially discovered at 3D structure level (20), and later analysis revealed significant sequence identity between the symmetry-related elements (9), allowing us to propose an evolutionary pathway for the modern ZIP fold (10). Taking advantage of the internal symmetry, we generated an outward-facing conformation model using repeat-swap homology modeling (23), and the proposed elevator motion was recapitulated by a later metadynamics simulation study (24). However, whether internal symmetry and sequence homology between the inverted repeats are common features in other ZIPs is still an open question.

Another prominent long-sought question is how different ZIP transporters exert their diverse biological functions. The answer likely lies in their distinct biochemical properties determined by subfamily- and type-specific features in sequence and 3D structure. Identifying these features and defining their roles are therefore essential for understanding functional specialization of ZIPs. Evolutionary approaches are often used to identify subfamily-specific features for later functional study. Among these, ancestral sequence reconstruction (ASR) has proven to be a powerful method for uncovering subgroup-specific characteristics by comparing reconstructed ancestral sequences with their extant successors (25,26). However, applying ASR to very large and ancient protein families remains challenging (27). In the case of the ZIP family, which diverged before the emergence of eukaryotic species, sequence identities between members of different subfamilies are often below 20%, complicating ancestral reconstruction.

In this study, by exploiting the internal symmetry broadly present in ZIPs, we generated ancestral sequences using ASR, revealing a significantly higher degree of sequence symmetry and thereby supporting the proposed evolutionary pathway of the modern ZIP fold. Leveraging this symmetrical structural framework, we identified a translocation pathway in the ancestral protein, which consists of symmetrical entry, transport, and exit sites, along with external and internal gates that separate them. Application of this structural framework to human ZIP4 enabled us to identify and characterize gating-forming residues and LIV-1 subfamily-specific residues, thereby advancing the understanding of metal transport by ZIP4. We anticipate that this method is applicable to other ZIP subfamilies and, more generally, to other ancient and highly divergent transporter families that retain internal symmetry.

## Results

### Internal symmetry is broadly present in prokaryotic ZIPs

The structure of BbZIP revealed two inverted repeats (**Figure 1A**): TM1-4 can be superimposed with TM5-8 when it is rotated by 180 degree along a pseudo two-fold axis that penetrates the structure center and is parallel to the membrane plane (20). Later, we reported that the amino acid sequences of the symmetry-related elements can also be aligned (9), indicating the homology between these symmetry-related elements. Further analysis led to the proposal that the current 8-TM ZIP fold evolved from a homodimer of a 4-TM ancestor that underwent gene duplication, domain rearrangement (circular permutation, **Figure S1**), and gene fusion (**Figure 1B**) (10). However, it is unclear whether the observed similarity at the sequence level represents a common feature among ZIPs or due to coincidence from random substitution.

**Figure 1.**
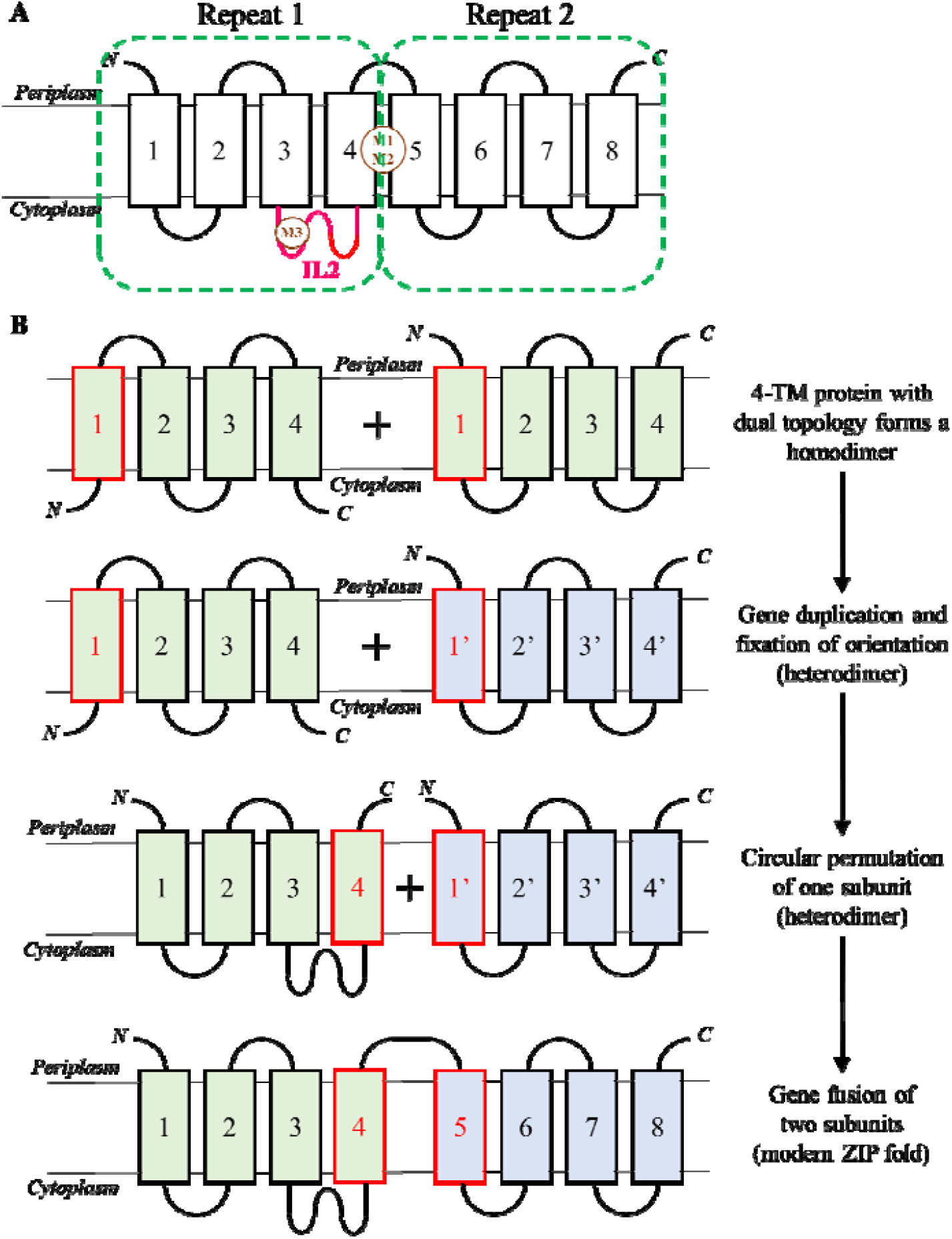
Proposed evolutionary pathway of the modern ZIP fold. (**A**) Topology of the modern ZIP fold. Transmembrane helices are shown as rectangles, and the inverted repeats are indicated by the dashed frames. Known metal binding sites, M1-M3, and the intracellular loop 2 (IL2, magenta) are labeled. (**B**) Evolution of the modern ZIP fold from an ancestral 4-TM protein. Figures are adapted from ref 10, licensed under CC BY 4.0. Illustration of circular permutation is shown in Figure S1.

In this work, we examined sequence similarity between the proposed symmetry-related elements, i.e. TM1 vs. TM6, TM2 vs. TM7, TM3 vs. TM8, and TM4 vs. TM5, in prokaryotic ZIPs from different subfamilies (**Figure 2**). As indicated in our previous sequence alignment and sequence similarity network analyses (9,10), the ZIP family consists of five major subfamilies, including GufA, LIV-1, ZIPI, ZIPII, and ZIP9, and many smaller groups. Among the major subfamilies, four of them (GufA, LIV-1, ZIPII, and ZIP9) contain members from prokaryotic and eukaryotic species, suggesting that divergence occurred before the emergence of eukaryotic ZIPs. We therefore chose two prokaryotic members from each subfamily and conducted sequence alignment. To increase the accuracy, we deployed 3D structure-guided sequence alignment using the structural models retrieved from AlphaFold Protein Structure Database. As shown in **Figure 2**, significant homology was detected between most symmetry-related TMs. Among the four pairs of TMs, the number of identical residues follows the order from large to small: TM2 vs. TM7, TM4 vs. TM5, TM1 vs. TM6, and TM3 vs. TM8. Notably, TM2/4/5/7 are the pore-forming TMs that are located in the inner circle of the structure, whereas TM1/3/7/8 form the peripheral circle. Thus, the pore-forming TMs show a higher degree of symmetry than the peripheral TMs. This analysis also revealed that the members in the GufA subfamily, including BbZIP and ZupT from *E*.*coli*, showed the highest sequence identity between the symmetry-related TMs.

**Figure 2.**
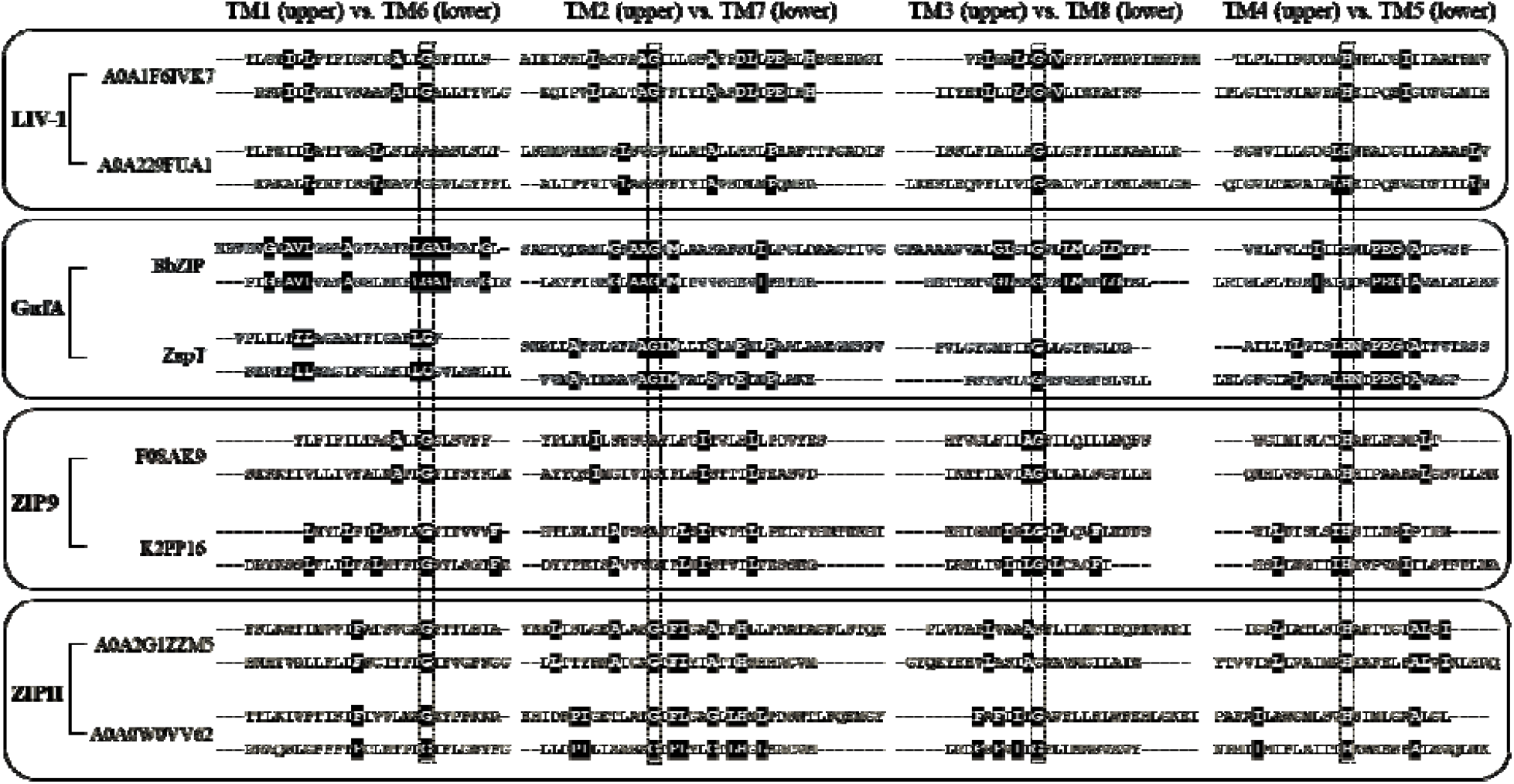
Detection of sequence identity between the proposed symmetry-related TMs in prokaryotic ZIPs from four ZIP subfamilies. In each subfamily (LIV-1, GufA, ZIP9, or ZIPII), two chosen sequences labeled with UniProt IDs were applied to structure-guided sequence alignment using the predicted structural models retrieved from AlphaFold Protein Structure Database and the BbZIP structure. The symmetry-related TMs were aligned based on the alignments of the inverted repeats in a BbZIP structure (PDB 5TSA). The residues identical between the proposed symmetry-related TMs are shown in white on a black background. Highly conserved residues highlighted in the dashed frames were used to gauge the alignments.

### Ancestral sequences showed a higher degree of internal symmetry

According to the proposed evolutionary pathway of the modern ZIP fold (**Figure 1B**), an extant ZIP with a higher degree of sequence identity between the symmetry-related TMs preserves a higher level of ancient features and thus is closer to the ancestral sequence by the time of gene duplication. Based on this consideration, we chose BbZIP and ZupT as input queries in ASR.

Using FireProt-ASR, a web server for automated ASR (28), we generated two sets of ancestral sequences derived from BbZIP and ZupT, respectively. Remarkably, for the root nodes in both ASR (BbZIP-derived Anc151 or BbAnc151 and ZupT-derived Anc151 or ZuAnc151), sequence alignment of the symmetry-related TMs showed a significantly increased sequence identity, particularly in the pore-forming TMs (TM2 vs. TM7, and TM4 vs. TM5), compared to their corresponding successors (**Figure 3A**). This result excludes the possibility that the sequence homology between symmetry-related TMs in selected ZIPs was a coincidence due to random substitutions, as the ancestral sequences calculated from hundreds of homologous sequences would otherwise show reduced, rather than increased, sequence identity between the proposed repeats. Instead, this result strongly supports the model that the modern 8-TM ZIP fold is evolved from an ancestral 4-TM protein (**Figure 1B**).

**Figure 3.**
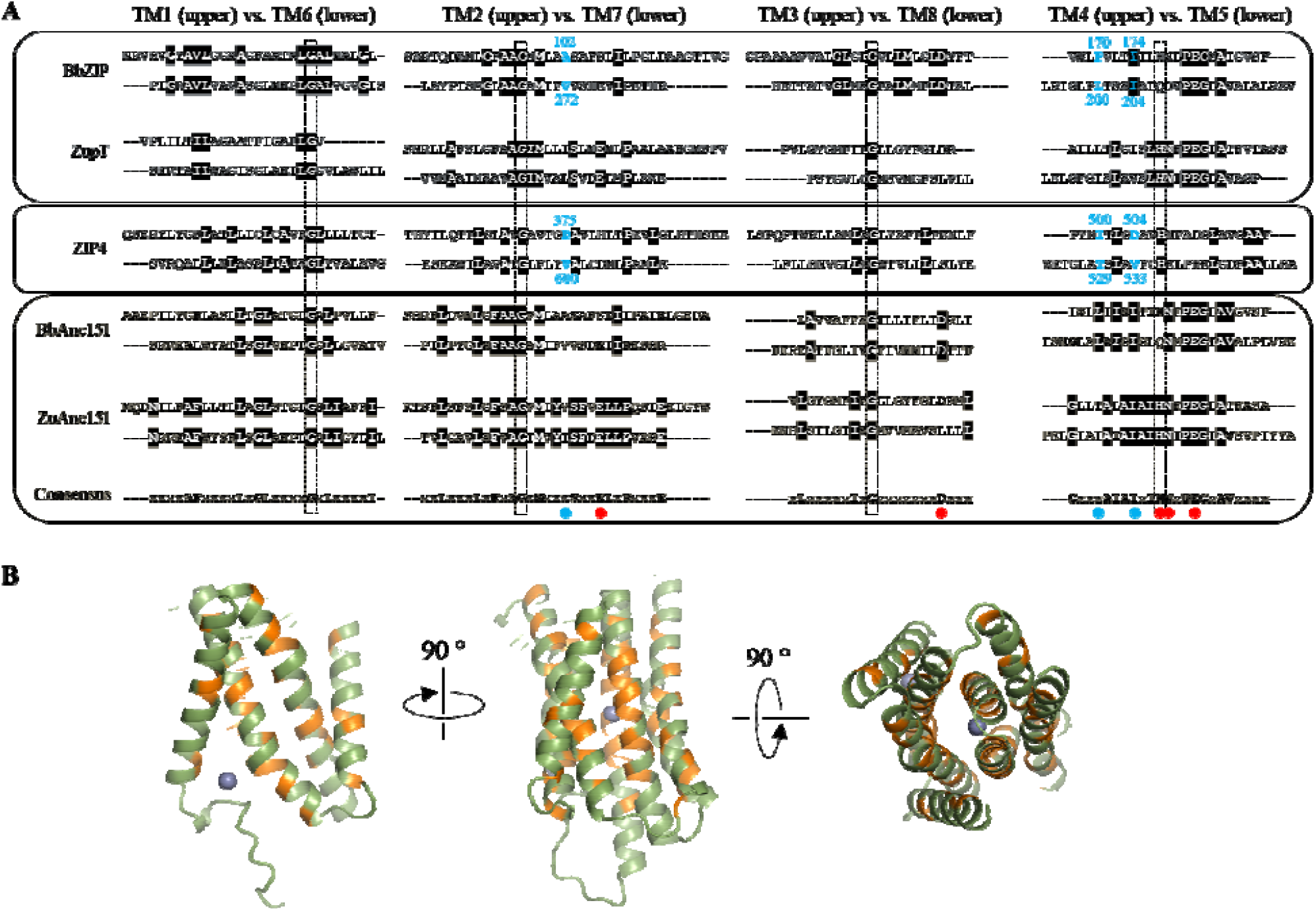
ASR for the ZIP family. (**A**) Comparison of the TM alignments between the successors (BbZIP and ZupT, light blue box) and the reconstructed ancestral sequences (yellow box). Only the root nodes (BbAnc151 and ZuAnc151, derived from BbZIP and ZupT, respectively) are shown. Highly conserved residues (in the dashed frames) were used to gauge the TM alignments. Only the residues that are identical at least in three of four ASR sequences are considered to be consensus and shown in bold in the consensus sequence. The metal chelating residues forming the entry, exit, and transport sites are marked by red dots. The positions where the gate-forming residues are located are marked by cyan dots and the corresponding residues in BbZIP and ZIP4 are labeled and colored in cyan. The alignment for the TMs of human ZIP4 (white box) is also included for later discussion. (**B**) Mapping the ancestral consensus residues on the structural model of ZuAnc151. The consensus residues are colored in orange. The grey spheres indicate zinc ions. The structural mode with two bound zinc ions was generated using AlphaFold 3.

Alignment of two root nodes (BbAnc151 and ZuAnc151) led to a consensus sequence that contains the residues likely present in the ancestral 4-TM protein at the time of gene duplication (**Figure 3A**). Mapping these residues on the AlphaFold 3 predicted structure model of ZuAnc151 revealed that most are located near the center of the structure (**Figure 3B**), indicating that ancestral sequence features have been preferentially preserved in the protein core, whereas the information at the peripheral regions has faded during evolution.

### Identification of a symmetrical metal translocation pathway

Labeling the metal chelating residues along the translocation pathway in ZuAnc151 revealed multiple metal binding sites (**Figure 4A**). In addition to the HNxPEG motif on TM4 and TM5 that form the transport site, a glutamate residue in TM7 (E223) joins an aspartate residue on TM3 (D84) to form a metal binding site at the cytoplasmic side. This site corresponds to the M3 site in BbZIP that is located at the end of the translocation pathway and thus functions as an exit site, playing a regulatory role by limiting metal release to the cytoplasm (22,24). Owing to the symmetrical architecture, a metal binding site with the same amino acid composition (E55 on TM2 and D252 on TM8) form an entry site of the translocation pathway, likely functioning to attract metal ions to enter the translocation pathway. Connecting these metal-binding sites defines a symmetrical metal relay pathway: metals enter through the entry site and reach the transport site by passing an open external gate while the internal gate remains closed; following the elevator motion of the transporter, metals are released from the transport site to the exit site through the now open internal gate, with the external gate closed to prevent backflow (**Figure 4A**).

**Figure 4.**
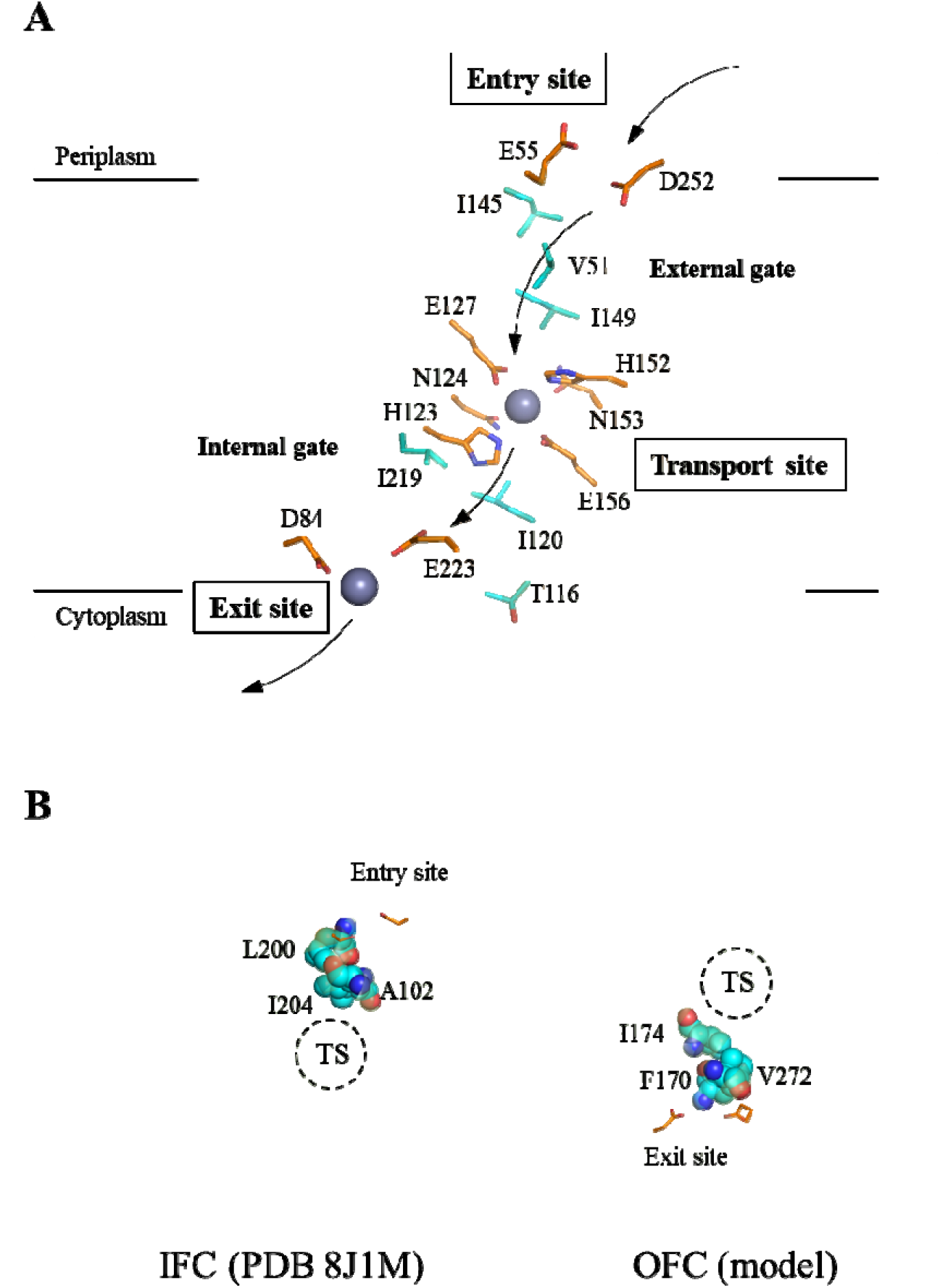
Symmetrical translocation pathway in the ancestral ZuAnc151. (**A**) Metal relay in the translocation pathway. The structure model of ZnAnc151 with bound zinc ions was generated by AlphaFold 3 and colored in the same manner as in Figure 3B. The residues involved in metal relay (entry, transport, and exit sites) and the gate-forming residues (cyan) are shown in stick mode. The arrows indicate how metal substrate translocate through the transporter. (**B**) Identification of the gate-forming residues in BbZIP. *Left*: External gate revealed in the IFC (PDB 8J1M). *Right*: Internal gate in an OFC model (ref 23). Both structures are in the same orientation as ZuAnc151 shown in (A). The gate-forming residues (cyan) are shown in sphere mode. The dashed circles indicate the transport sites (TS).

To identify gate-forming residues in ZuAnc151, we leveraged the internal symmetry of the BbZIP structure. In the experimentally solved IFC of BbZIP, A102 (TM2), L200 and I204 (TM5) block the passage of metal from the entry site to the transport site, thereby forming the external gate (**Figure 4B**). Taking advantage of structural symmetry (**Figures 2** and **3A**), residues symmetrically related to those forming the external gate are expected to constitute the internal gate. In BbZIP, these residues, including V272 (TM7), F170 and I174 (TM4) (**Figure 3A**), indeed block the pathway from the transport site to the exit site in the OFC model that was built using repeat-swap homology modeling in our previous study (**Figure 4C**) (23). Correspondingly, the gate-forming residues in ZuAnc151 (V51, I145, and I149 for the external gate and T116, I120, and I219 for the internal gate) were identified through sequence alignment (**Figure 3A**) and mapped onto the metal translocation pathway (**Figure 4**). Notably, most of these residues are hydrophobic, suggesting that ancestral ZIPs likely use hydrophobic seals to prevent metal leakage to maintain alternating access.

### Functional study of non-metal chelating residues along the translocation pathway of human ZIP4

Next, we applied the insights gained from the above analysis to human ZIP4, a representative eukaryotic ZIP with well-established functional assays (29-32). We first identified the gate-forming residues in ZIP4 from the sequence alignment with BbZIP, in which the gate-forming residues have been determined using experimentally solved structures and symmetry-based analysis (**Figures 3A** and **4**). Accordingly, D375 (TM2), and T529 and V533 (TM5) form the external gate in the IFC, whereas I500 and D504 (TM4), and V600 (TM7) form the internal gate in the OFC (**Figures 3A**). Using ColabFold with optimized subsampling parameters for multiple sequence alignment (33), we generated ZIP4 structural models in distinct conformational states. Inspection of the predicted IFC and outward-facing conformation (OFC) models confirmed that the proposed gates separate the entry and exit sites from the transport site (**Figure 5A**), respectively.

**Figure 5.**
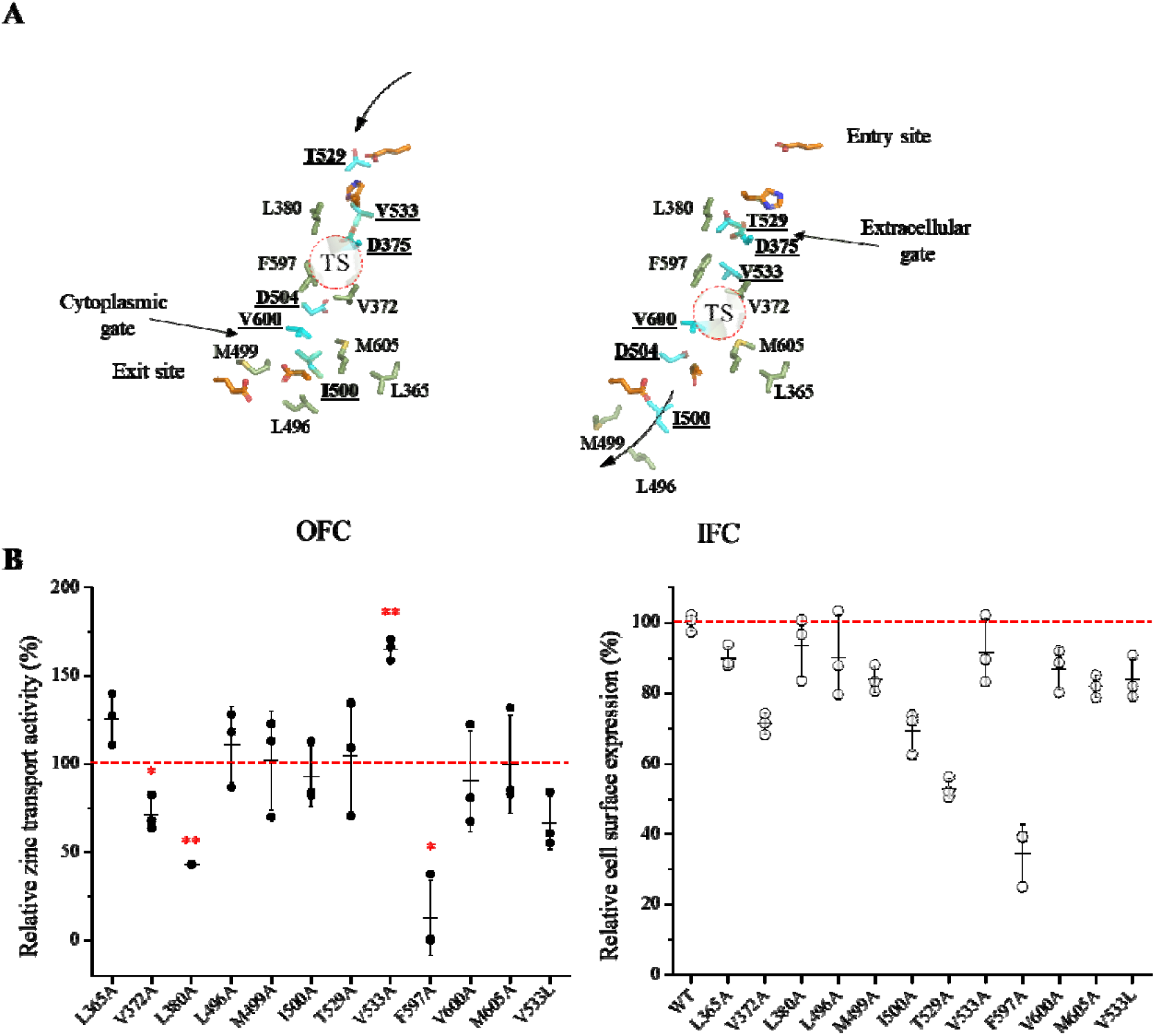
Identification and characterization of the gate-forming residues in human ZIP4. (**A**) Metal translocation pathway in the OFC and IFC models, which were generated by ColabFold via subsampling in multiple sequence alignment. The gate-forming residues (cyan and underlined) and the other hydrophobic residues along the pathway (light green) are shown in stick mode. The residues forming the entry site or exit site are colored in orange. The dashed circles indicate the transport site (TS). The arrows indicate where metal substrate enters and leaves the translocation pathway in the OFC and IFC states, respectively. (**B**) Mutagenesis and functional characterization of the residues along the translocation pathway. *Left*: zinc transport activities of the variants. The data shown are from three independent experiments with each data point representing the mean from one experiment. Three biological replicates were included in each experiment. The activities are expressed as the percentages of that of wild-type ZIP4, and the red dashed line indicates 100% activity. One-sample two-tailed *t* tests were conducted to examine the null hypothesis. The *P* values for V372A, L380A, V533A, and F597A are 0.034, 0.0032, 0.003, and 0.019, respectively. ^*^, *P*<0.05; ^**^, *P*<0.01. *Right*: Cell surface expression levels of the variants determined by flow cytometry. The data shown are from one of two independent experiments with similar results. An Alexa Fluor 647-conjugated anti-HA monoclonal antibody was used to label ZIP4 variants expressed on the surface of live cells. Cell surface expression levels of the ZIP4 variants are expressed as the percentages of that of wild-type ZIP4.

In contrast to the ancestral sequences, where gate-forming residues are predominantly hydrophobic, both the external and internal gates of ZIP4 each contain an aspartate residue. However, kinetic studies of ZIP4 do not support leaky gates that would otherwise result in channel-like activity (30,31,34-36), suggesting that ZIP4 may still rely on hydrophobic residues to prevent gate leakage. We therefore examined the non-metal chelating residues in the ZIP4 gates to probe their roles in controlling metal transport. T529, V533, I500, and V600, were individually substituted with alanine and their zinc transport activities were measured using the cell-based ^70^Zn transport assay (**Figure 5B**) (32). I500A, T529A, and V600A did not affect activity, whereas V533A led to a significantly increased activity by ∼60%. Flow cytometry analysis revealed no significant changes in cell surface expression for V533A, I500A, and V600A; however, T529A exhibited a ∼50% reduction in surface expression. After normalization to surface expression, T529A showed an increased intrinsic transport activity. Substitution of V533 with a bulkier leucine residue (V533L) reduced activity by ∼30%, but this reduction was not statistically significant (*P*=0.063). Together, these results indicate that T529 and V533 in the external gate play key roles in regulating metal flux, whereas residues forming the internal gate appear less critical.

To determine whether additional hydrophobic residues within the translocation pathway contribute to transport regulation, we performed alanine scanning on L365, V372, L380, L496, M499, F597, and M605 based on ZIP4 structural models (**Figure 5A**). As shown in **Figure 5B**, while most substitutions had little effect on activity, V372A and L380A moderately reduced transport, and F597A caused a drastic reduction of more than 80%. The reduced activities of V372A and F597A are partially attributable to decreased cell surface expression. None of these variants exhibited increased activity, in contrast to T529A and V533A, suggesting that they play roles distinct from residues in the external gate.

Remarkably, all mutations that altered transport activity (L372A, L380A, T529A, V533A, and F597A) cluster at or near the external gate, whereas residues proximal to the internal gate are more tolerant to mutagenesis, suggesting that the structural integrity of the external gate is crucial in controlling metal translocation. This asymmetry is unexpected given the overall structural symmetry of the ZIP fold, and may reflect evolutionary specialization from an ancestral bidirectional transporter toward a functionally unidirectional system.

### Identification and functional characterization of the LIV-1 unique residues

The LIV-1 subfamily represents one of the largest groups in the ZIP family, comprising nine of the fourteen human ZIP transporters. A defining feature of this subfamily is a signature motif located on TM5, HExxHExxD, which was recognized since the discovery of LIV-1 proteins (37,38). Comparison with the consensus ancestral sequence revealed that the first three conserved residues in this motif are either identical to the ancestor sequence (H536) or arise from conservative substitutions (E537 and H540) (**Figure 6A**). In contrast, E541 and D544 correspond to glycine and valine, respectively, in the ancestral sequence. Additionally, sequence analysis identified an aspartate residue (D504) in TM4, also a gate-forming residue, that replaces an isoleucine in the ancestral sequence. Notably, D504, E541, and D544 are spatially proximal in the predicted structural model of ZIP4 (**Figure 6B**), implying a potential functional relationship. To investigate their roles in zinc transport, we generated alanine substitutions at each position and performed metal transport assays. As shown in **Figure 6C**, the E541A mutation drastically reduced transport activity, whereas D504A and D544A significantly increased activity by 2-3 folds. Flow cytometry analysis revealed that the D504A variant exhibits a moderate reduction in cell surface expression (**Figure 6C**). Given its increased transport activity, this mutation results in a marked enhancement of transport efficiency. In contrast, the substantial reduction in activity observed for E541A is not accompanied by decreased surface expression, indicating that this variant has severely impaired transport efficiency.

**Figure 6.**
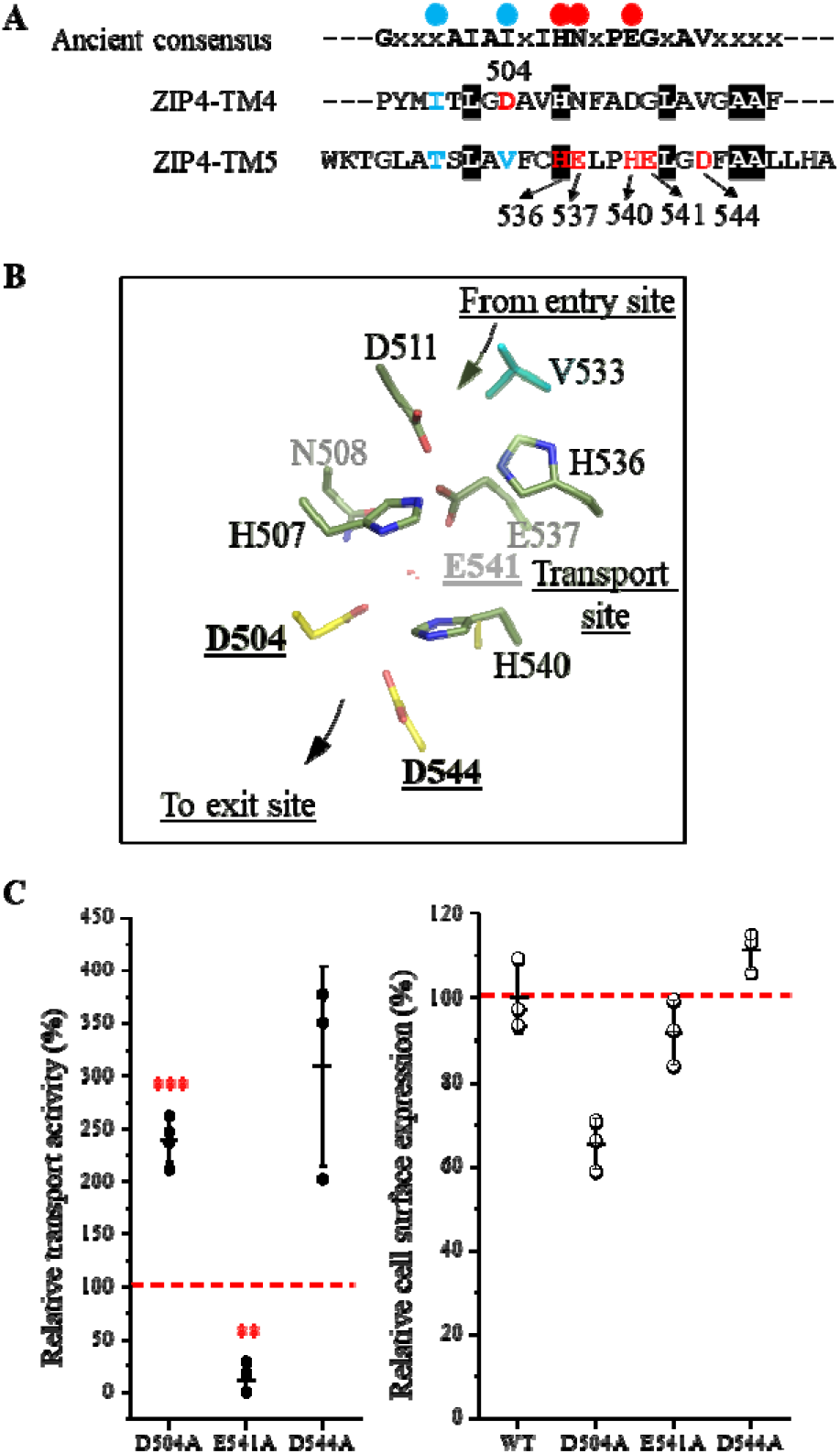
Characterization of the LIV-1 unique metal-chelating residues on TM4 and TM5. (**A**) Sequence comparison of TM4 and TM5 of ZIP4 with the consensus sequence of ancestral proteins (Figure 3A). The LIV-1 signature motif and D504 are labeled and colored in red. The red and blue dots indicate the residues contributive to the transport site and the gates, respectively. (**B**) Structure of D504, E541, and D544 in the ZIP4 model in the IFC. The residues contributing to the transport site (pale green), the external gate (cyan), and D504, E541, and D544 (yellow) are shown in stick mode. (**C**) Functional characterization of D504A, E541A, and D544A. *Left*: zinc transport activities of the variants. The activities are expressed as the percentages of that of wild-type ZIP4, and the red dashed line indicates 100% relative activity. The data shown are from three to four independent experiments with each data point representing the mean of one experiment. Three biological replicates were included in each experiment. One-sample *t* test was conducted to examine the null hypothesis. The *P* values for D504A, E541A, and D544A are 0.0009, 0.0609, and 0.0011, respectively. ^**^, *P*<0.01; ^***^, *P*<0.001. *Right*: Cell surface expression levels of the variants determined by flow cytometry. The data shown are from one of two independent experiments with similar results. An Alexa Fluor 647-conjugated anti-HA monoclonal antibody was used to label ZIP4 variants expressed on the surface of live cells. Cell surface expression levels of the ZIP4 variants are expressed as the percentages of that of wild-type ZIP4.

In the IFC model of ZIP4, D504 and D544 are positioned in close proximity to H540, a key residue contributing to the transport site. In our previous study, we demonstrated that the H540A mutation increases transport activity by more than twofold (31). Subsequent structural and computational studies suggest that H540 plays a critical role in metal release by flipping its side chain to facilitate this process (24). Loss of the side chain may therefore enhance metal release and increase transport rates. If D504 and D544 form a metal binding site together with H540, as suggested by the structural model (**Figure 6B**), disruption of this site, which is immediately downstream of the transport site, would be expected to facilitate metal release and thereby enhance transport. E541 is also located near this region; however, the severe loss of activity observed for E541A suggests that this residue may serve additional roles. For example, E541 may play a structural role to stabilize the local structure near the transport site by participating in the extensive hydrogen-bond network. Further elucidation of the mechanisms underlying these activity changes will require further structural, biochemical, and computational studies.

## Discussion

Despite the critical role of ZIP transporters in maintaining trace element homeostasis, the mechanisms by which they translocate metal substrates remain elusive. This knowledge gap largely stems from the limited availability of experimentally solved ZIP structures in distinct functional states. In this study, we leveraged the internal symmetry of the ZIP fold to gain new insights into the movement of metal substrates along the translocation pathway.

The presence of inverted repeats is a hallmark of many membrane transporters and reflects their evolutionary pathway through gene duplication and fusion events. In this work, we demonstrate that the internal symmetry observed in BbZIP extends beyond this prototypical member. Analysis across major ZIP subfamilies revealed that sequence homology between the proposed inverted repeats is broadly present among prokaryotic ZIPs (**Figure 2**).

Notably, members of the GufA subfamily exhibit the highest degree of sequence identity, suggesting that ZIPs such as BbZIP and ZupT retain more ancestral features than members of other subfamilies. ASR of an ancient and highly diverse family such as ZIP is inherently challenging because low sequence conservation obscures the evolutionary signal needed to reliably infer ancestral features. Exploiting the internal symmetry of ZIPs, particularly the symmetry retained at the amino acid sequence level, provides an independent structural and evolutionary constraint that helps recover these features. Consistent with this idea, using BbZIP and ZupT as input queries, ASR yielded ancestral sequences with even greater sequence identity between the repeats (**Figure 3**). These findings strongly support the hypothesis that the modern ZIP fold arose through gene duplication and fusion of an ancestral 4-TM protein (**Figure 1**).

The structural model of one of the reconstructed ancestral sequences (ZuAns151) enabled the identification of a highly symmetrical metal translocation pathway (**Figure 4**). This pathway is defined by symmetrically positioned metal-chelating residues that form the entrance and exit sites. The central transport site is primarily composed of metal chelating residues from TM4 and TM5, which themselves are symmetry-related. Flanking this site are the external and internal gates, which are also arranged symmetrically to separate the transport site from the entry site in the IFC and from the exit site in the OFC, respectively.

Exploiting this symmetry framework, we identified the residues forming external or internal gates of ZIP4 (**Figure 5**). Functional characterization revealed that two residues, T529 and V533, located within the external gate, play a critical role in controlling transport. Comparison of ZIP4 with the ancestral sequences revealed three metal-chelating residues along the translocation pathway that appear to have emerged later in evolution (**Figure 6**). Two of these residues, E541 and D544, are within the conserved LIV-1 signature motif (HExxHExxD). Functional analyses demonstrated that E541 is essential for zinc transport, whereas alanine substitutions at D504 and D544 enhance transport activity. Structural modeling suggests that D504, D544, and the neighboring H540 may form a metal-binding site immediately downstream of the transport site toward the exit (**Figure 6B**), potentially slowing metal release by transiently retaining metal within the translocation pathway. This mechanism is reminiscent of the second intracellular loop (IL2) of BbZIP and ZIP4, where replacement of clustered histidine residues leads to increased transport rates (24).

Collectively, our results demonstrate that integrating symmetry analysis with ASR provides a unique perspective for elucidating the working mechanism conserved in the entire transporter family and enables the identification of elements that emerged in later evolution to confer functional specialization and regulatory control. Application of this approach to the ZIP family is expected to reveal additional subfamily- and type-specific features that determine distinct transport properties and regulatory mechanisms underlying their non-redundant biological functions.

## Methods

### Gene, plasmid, and Cell line

The complementary DNA of human ZIP4 (GenBank ID: BC062625) from Mammalian Gene Collection was purchased from GE Healthcare. The DNA encoding ZIP4 were inserted into a modified pEGFP-N1 vector (Clone, Cat#6085-1) in which the downstream EGFP gene was deleted and a human influenza hemagglutinin (HA) tag was inserted before the stop codon, leading to a ZIP4-HA construct. This vector was further modified by inserting a FLAG immediately after the signal peptide for a double-tagged construct (FLAG-ZIP4-HA). The variants shown in Figure 5 were double-tagged, whereas those in Figure 6 are HA-tagged. Plasmids were purified using Miniprep (Promega, cat#A1460). All the primer sequences for mutagenesis are listed in **Table S1**.

Human embryonic kidney cells (HEK293T, ATCC, Cat#CRL-3216) were cultured in Dulbecco’s Modified Eagle Medium (DMEM, Thermo Fisher Scientific, Invitrogen, Cat#11965092) supplemented with 10% (v/v) fetal bovine serum (Thermo Fisher Scientific, Invitrogen, Cat#15240062) at 5% CO_2_ and 37°C.

### Zinc transport assay and inductively coupled plasma mass spectrometry (ICP-MS)

For transient expression of ZIP4 and its variants, HEK293T cells were transfected with the corresponding plasmids using Lipofectamine 2000 (Thermo Fisher Scientific, Invitrogen, Cat#11668019) and cultured for 24 h prior to the transport assay.

^70^Zn transport assay and ICP-MS measurement were conducted as described (32). In brief, 24 h post transfection, HEK293T cells expressing ZIP4 or variants were washed with a washing buffer (10 mM HEPES, 142 mM NaCl, 5 mM KCl, 10 mM glucose, pH 7.3). 5 μM ^70^Zn (American Elements, Product# ZN-OX-01-ISO.070I) was added to cells to initiate zinc transport. After incubation at 37°C for 30 min, cells were transferred on ice and an equal volume of the ice-cold washing buffer containing 1 mM EDTA was added to the cells to terminate metal uptake. After three washes with the ice-cold washing buffer, 70% trace nitric acid was added to cells for digestion and subsequent ICP-MS analysis. Zinc uptake by cells was expressed as the count ratio of ^70^Zn/^31^P determined by ICP-MS, and zinc transport activity was calculated by subtracting zinc uptake by the cells transfected with an empty vector from that by the cells expressing metal transporters.

### Cell surface expression level of ZIP4 determined by flow cytometry

To determine and compare the cell surface expression levels of ZIP4 and its variants, HEK293T cells transiently transfected with the plasmids encoding ZIP4 or its variants were trypsinized, washed with prechilled DPBS without Ca^2+^ and Mg^2+^ (Gibco, Cat# 14190094), and put on ice for 15 min to quench transporter trafficking. Cells were then incubated with Alexa Fluor 647-conjugated anti-HA monoclonal antibody (Invitrogen, Cat# 26183-A647) diluted 1:400 at 4 °C with gentle agitation for 1 h. After washing with DPBS without Ca^2+^ and Mg^2+^, cells were fixed with 4% formaldehyde at room temperature for 10 min and then washed and resuspended in DPBS.

Cell samples were applied to a ThermoFisher Attune Cytpix flow cytometer. At least 10,000 cells were assessed for each sample. Cell debris was excluded based on FSC-A vs. SSC-A panel and single cells were selected through FSC-A vs. FSC-H gating (**Figure S2**). Mean fluorescence intensities (MFIs) of Alexa Fluor 647 were calculated for each sample. All the flow cytometry data processing was performed in FlowJo 10.8.1.

### Ancestral sequence reconstruction

Ancestral sequence reconstruction was performed using the FireProt-ASR web server (https://loschmidt.chemi.muni.cz/fireprotasr/). The sequences of BbZIP (UniProt ID: A0A0H3LM39) and ZupT from *E. coli* (UniProt ID: P0A8H3) served as the query inputs. Calculations were conducted using default parameters, with the exception of specifying conserved metal-chelating residues within the transport site as essential residues to ensure the functional relevance and quality of the retrieved homologs. For each protein, a dataset of 150 homologous sequences was retrieved and used to reconstruct the ancestral nodes.

### Structure prediction

AlphaFold 3 (https://alphafoldserver.com/) was used to predict the structures of prokaryotic ZIPs whose structures have not been experimentally determined, which were used to guide sequence alignment. AF3 was also used to predict the structure of ancestral proteins generated by ASR with bound zinc ions (**Figures 3B** and **4**).

ColabFold (https://colab.research.google.com/github/sokrypton/ColabFold/blob/main/AlphaFold2.ipynb) was used to predict structures of human ZIP4 (UniProt ID: Q6P5W5). To facilitate the generation of distinct conformational states, the parameters for multiple sequence alignment sampling were chosen as below: max_msa=32:64, num_seeds=8, and num_recycles=5. The pLDDT values of the transmembrane domains of the OFC and IFC models shown in **Figure 5** are 78.3 and 83.2, respectively.

## Quantification and statistical analysis

We assumed a normal distribution of the samples and multiple comparisons were assessed using One-sample two-tailed Student’s *t-*test. A *P*< 0.05 was considered statistically significant. Data were presented as mean ± standard deviation.

## Data availability statement

Further information and requests for resources and reagents should be directed to Dr. Jian Hu. Any additional information required to reanalyze the data reported in this paper is available from Dr. Jian Hu upon request.

## Acknowledgement

The data presented herein were obtained using instrumentation in the MSU Flow Cytometry Core Facility. The facility is funded in part through the financial support of Michigan State University’s Office of Research & Innovation and Colleges of Osteopathic Medicine, Human Medicine, Veterinary Medicine, Natural Sciences, and Engineering. The Attune CytPix is supported by the Equipment Grants Program, award #2022-70410-38419, from the U.S. Department of Agriculture, National Institute of Food and Agriculture. We thank Quantitative Bio Element Analysis and Mapping (QBEAM) center at Michigan State University for the assistance of using ICP-MS. We thank Dr. Yuhan Jiang and Michael Nikolovski at Michigan State University for running ICP-MS experiments. This work was supported by NIH grant GM140931 (to J.H.).

## Author contributions

Investigation: Y.Z., T.W., and Z.H. conducted biochemical experiments and data analysis, and J.H. conducted bioinformatic analysis; Writing: Y.Z., T.W., Z.H., and J.H.; Conceptualization: J.H.; Supervision: J.H.

