## Supplementary Information for "Symmetry Analysis and Ancestral Sequence Reconstruction Reveal a Symmetrical Translocation Pathway and Activity Determinants of ZIP Metal Transporter"

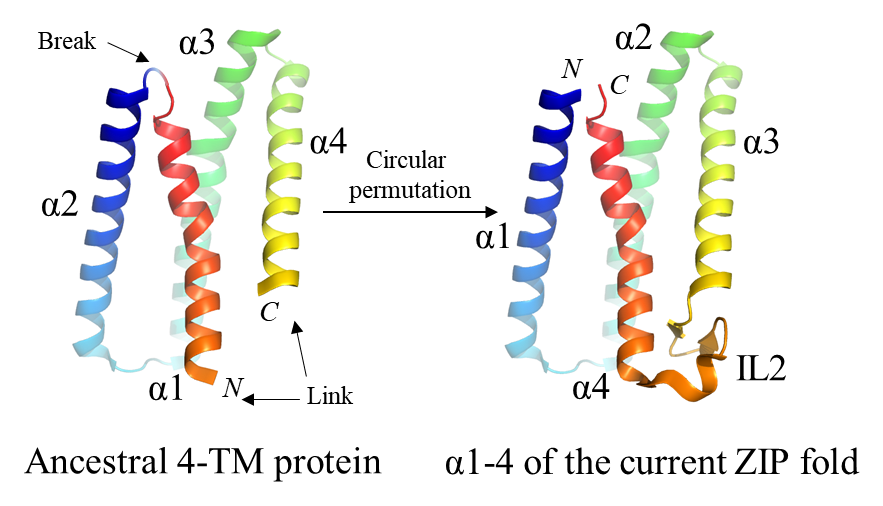

**Figure S1.** Illustration of the proposed circular permutation. TM1-4 in the modern ZIP fold is derived from an ancestral 4-TM protein through circular permutation, which reorders the TMs without affecting the packing of TMs in the 3D structure.

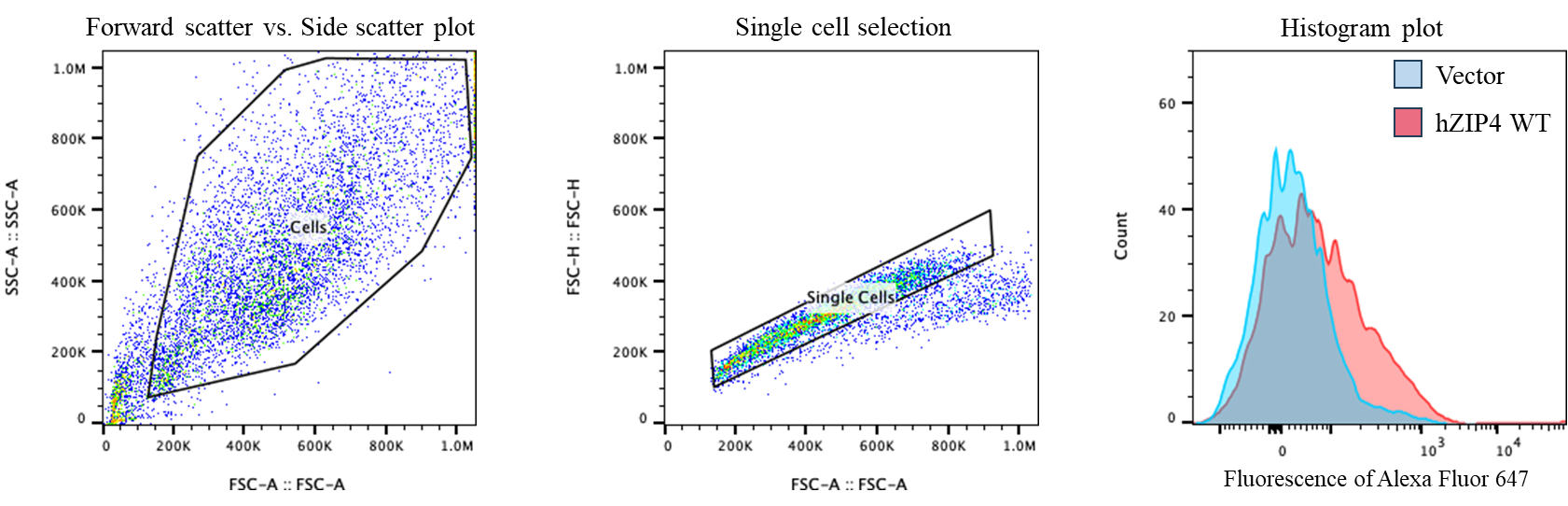

**Figure S2.** Flow chart for flow cytometry data processing.

**Table S1.** The primers used in this study.

| Primer | Sequence (forward only, 5’-3’) |
| --- | --- |
| L365A | CACTACATCCTGCAGACCTTCGCGAGCCTGGCAGTGGGTGCACTC |
| L372A | CTGAGCCTGGCAGTGGGTGCAGCGACTGGGGACGCTGTCCTGCAT |
| L380A | ACTGGGGACGCTGTCCTGCATGCGACGCCCAAGGTGCTGGGGCTG |
| L496A | CTGAGCCCAGAGTTGAGGCTAGCGCCCTATATGATCACTCTGGGC |
| M499A | GAGTTGAGGCTACTGCCCTATGCGATCACTCTGGGCGACGCCGTG |
| I500A | TTGAGGCTACTGCCCTATATGGCGACTCTGGGCGACGCCGTGCAC |
| T529A | TCCTGGAAGACCGGGCTGGCCGCGTCGCTGGCCGTGTTCTGCCAC |
| V533A | GGGCTGGCCACCTCGCTGGCCGCGTTCTGCCACGAGTTGCCACAC |
| F597A | CTGGCAGTGGCCACCGGCCTGGCGCTCTACGTAGCACTCTGCGAC |
| V600A | GCCACCGGCCTGTTCCTCTACGCGGCACTCTGCGACATGCTCCCG |
| M605A | CTCTACGTAGCACTCTGCGACGCGCTCCCGGCGATGTTGAAAGTA |
| V533L | GGGCTGGCCACCTCGCTGGCCCTGTTCTGCCACGAGTTGCCACAC |
| D504A | CCCTATATGATCACTCTGGGCGCCGCCGTGCACAACTTCGCCGAC |
| E541A | GCCGTGTTCTGCCACGAGTTGCCACACGCCCTGGGGGACTTCGCCGCCTTGCTGCAC |
| D544A | GAGTTGCCACACGAGCTGGGGGCCTTCGCCGCCTTGCTGCACGCG |
